# Somatosensory processing deficits and altered cortico-hippocampal connectivity in *Shank3b^−/−^* mice

**DOI:** 10.1101/2021.02.28.433258

**Authors:** Luigi Balasco, Marco Pagani, Luca Pangrazzi, Evgenia Schlosman, Lorenzo Mattioni, Alberto Galbusera, Giovanni Provenzano, Alessandro Gozzi, Yuri Bozzi

**Author notes:** **Corresponding author:** Prof. Yuri Bozzi, **Email:**. **Author Contributions:** LB designed and performed behavioral and *in situ* hybridization experiments, analyzed data, and wrote the manuscript; MP performed rsfMRI experiments, analyzed data, and drafted the paper; LP, ES, and LM performed behavioral experiments; AGa provided technical assistance for rsfMRI experiments; GP, performed behavioral experiments and edited the manuscript; AGo, designed rsfMRI experiments, analyzed data, provided funding, and edited the manuscript; YB performed *in situ* hybridization experiments, analyzed data, supervised the whole study, provided funding, and wrote the manuscript. **Competing Interest Statement:** The authors declare no competing interests.

## Abstract

Abnormal tactile response is considered an integral feature of Autism Spectrum Disorders (ASDs), and hypo-responsiveness to tactile stimuli is often associated with the severity of ASDs core symptoms. Patients with Phelan-McDermid syndrome (PMS), caused by mutations in the SHANK3 gene, show ASD-like symptoms associated with aberrant tactile responses. However, the neural underpinnings of these somatosensory abnormalities are still poorly understood. Here we investigated, in *Shank3b^−/−^* adult mice, the neural substrates of whisker-guided behaviors, a key component of rodents’ interaction with the surrounding environment. To this aim, we assessed whisker-dependent behaviors in *Shank3b^−/−^* adult mice and age-matched controls, using the textured novel object recognition (tNORT) and whisker nuisance (WN) test. *Shank3b^−/−^* mice showed deficits in whisker-dependent texture discrimination in tNORT and behavioral hypo-responsiveness to repetitive whisker stimulation in WN. Notably, sensory hypo-responsiveness was accompanied by a significantly reduced activation of the primary somatosensory cortex (S1) and hippocampus, as measured by c-*fos* mRNA *in situ* hybridization, a proxy of neuronal activity following whisker stimulation. Moreover, resting-state fMRI showed a significantly reduced S1-hippocampal connectivity in *Shank3b* mutant mice. Together, these findings suggest that impaired crosstalk between hippocampus and S1 might underlie *Shank3b^−/−^* hypo-reactivity to whisker-dependent cues, highlighting a potentially generalizable form of dysfunctional somatosensory processing in ASD.

**Significance Statement:** Patients with Phelan-McDermid syndrome, a syndromic form of ASD caused by mutation of the SHANK3 gene, often show aberrant responses to touch. However, the neural basis of atypical sensory responses in ASD remains undetermined. Here we used *Shank3* deficient mice to investigate the neural substrates of behavioral responses to repetitive stimulation of the whiskers, a highly developed sensory organ in mice. We found that mice lacking the *Shank3* gene are hypo-responsive to repetitive whisker stimulation. This trait was associated with reduced engagement and connectivity between the primary somatosensory cortex and hippocampus. These results suggest that dysfunctional cortico-hippocampal coupling may underlie somatosensory processing deficits in SHANK3 mutation carriers and related syndromic forms of ASD.

## Introduction

Autism spectrum disorders (ASDs) represent a heterogeneous group of neurodevelopmental disorders characterized by social interactions and communication deficits, accompanied by restricted and stereotyped behaviors (1). Several studies also indicate that abnormal sensory processing is a crucial feature of ASD. About 90% of ASD individuals have atypical responses to different types of sensory stimuli (2), and sensory abnormalities (described as both hyper- or hypo-reactivity to sensory stimulation) are currently recognized as diagnostic criteria of ASD (1). Several pieces of evidence support the idea that altered sensory perception underlies ASD patients’ ability to interact with the surrounding environment, especially in the presence of novel stimuli and unexpected context (2). Both over- and under-responsiveness to tactile stimuli are frequently observed in ASD and fall under the general term of tactile defensiveness (3, 4). Most importantly, abnormal responses to tactile stimuli correlate with and predict the severity of ASD. Hypo-responsiveness to tactile stimuli is associated with higher severity of ASD core symptoms (5), and touch avoidance during infancy is predictive of ASD diagnosis in toddlers (6).

Phelan-McDermid syndrome (PMS) is a neurodevelopmental disorder characterized by ASD-like behaviors, developmental delay, intellectual disability, and absent or severely delayed speech (7). PMS is caused by mutations in the SHANK3 gene coding for SH3 and multiple ankyrin repeat domains protein 3 (8, 9), a crucial component of excitatory postsynaptic density (10, 11). Patients with PMS often show somatosensory processing dysfunction, including over-sensitiveness to touch and tactile defensiveness (12).

Altered somatosensory responses have been recently described in mouse strains harboring mutations in ASD-relevant genes, suggesting that whisker-dependent behaviors are a good proxy to study somatosensory processing defects relevant for ASD (4, 13). Mice use their whiskers to explore objects (14) and interact with conspecifics (15), thus allowing a detailed representation of the surrounding environment during navigation (16, 17).

Mice lacking the *Shank3* gene are widely considered as a reliable model to study ASD-like symptoms relevant to PMS (18, 19). Interestingly, *Shank3b^−/−^* mutant mice display aberrant whisker-independent texture discrimination and over-reactivity to tactile stimuli applied to hairy skin (20). More recent findings showed that developmental loss of *Shank3* in peripheral somatosensory neurons (that causes over-reactivity to tactile stimuli) also results in ASD-like behaviors in adulthood (21). While deficits of peripheral somatosensory neurons result in aberrant tactile responses and ASD-relevant behaviors in *Shank3* mutants, the neural substrates of whisker-dependent behaviors have been poorly investigated in this model.

Based on somatosensory processing deficits reported in PMS patients (12), we hypothesized that similar abnormalities were present in *Shank3b^−/−^* mice. Thus, we investigated the neural substrates of somatosensory responses in adult *Shank3b^−/−^* and control mice. *Shank3b^−/−^* mice showed hypo-reactivity to whisker-dependent cues, accompanied by reduced expression of the immediate-early gene c-*fos* in the primary somatosensory cortex (S1) and hippocampus. Finally, we used rsfMRI to probe functional connectivity between cortico-hippocampal regions that exhibited a reduced c-*fos* response. Our results indicate that *Shank3b^−/−^* mice show hypo-connectivity within the somatosensory-hippocampal network.

## Results

### *Shank3b^−/−^* mice show aberrant texture discrimination through whiskers

Human research suggests that PMS is associated with aberrant sensory responses. To test the presence of a similar dysfunction in *Shank3* mutant mice, we first tested *Shank3b^−/−^* and control mice in a whisker-dependent version of tNORT (22) (Fig. 1*A*) using sandpaper-wrapped cylinders that differ only in texture (smooth or rough; see Materials and Methods). As a general measure of locomotor activity and generalized anxiety we first assessed *Shank3b^−/−^* and control mice in an open field arena in two consecutive days. During both days of testing, *Shank3b^−/−^* mice showed a significantly reduced distance travelled (Fig. 1*B;* two-way ANOVA, *Shank3b^+/+^* vs. *Shank3b^−/−^*; main effect of genotype F(_1, 34)_ = 103.7; p<0.0001; main effect of testing days F_(1, 34)_ = 99.42, p<0,0001; post hoc Tukey’s test, *Shank3b^+/+^* vs. *Shank3b^−/−^* within day 1 and 2, p<0.0001) and time spent moving (*SI Appendix*, Fig. S1) compared to control littermates. Both genotypes showed a significant reduction of distance travelled (Fig. 1*B;* Tukey’s post hoc following two-way ANOVA, *Shank3b^+/+^* day 1 vs. *Shank3b^+/+^* day 2 and *Shank3b^−/−^* day 1 vs. *Shank3b^−/−^* day 2; p<0.0001) and time spent moving (*SI Appendix*, Fig. S1) between testing days, indicating habituation to the novel environment. Both genotypes also spent a comparable amount of time in center and borders of the open field arena (Fig. 1*C*) over the two days of habituation (three-way ANOVA, *Shank3b^+/+^* vs. *Shank3b^−/−^*; main effect of genotype F_(1, 68)_ = 0.03371, p=0.8549; main effect of days F_(1, 68)_ = 0.03002, p=0.8630; main effect of arena regions F_(1, 68)_ = 2346, p<0,0001). Finally, both *Shank3b^−/−^* and control mice displayed a preference for borders regions of the arena during both days of testing (Fig. 1*C*; Tukey’s post hoc following three-way ANOVA, center vs. borders within *Shank3b^+/+^* and center vs. borders within *Shank3b^−/−^*; p<0.0001), indicating a similar anxious behavior in both genotypes.

**Fig. 1.**
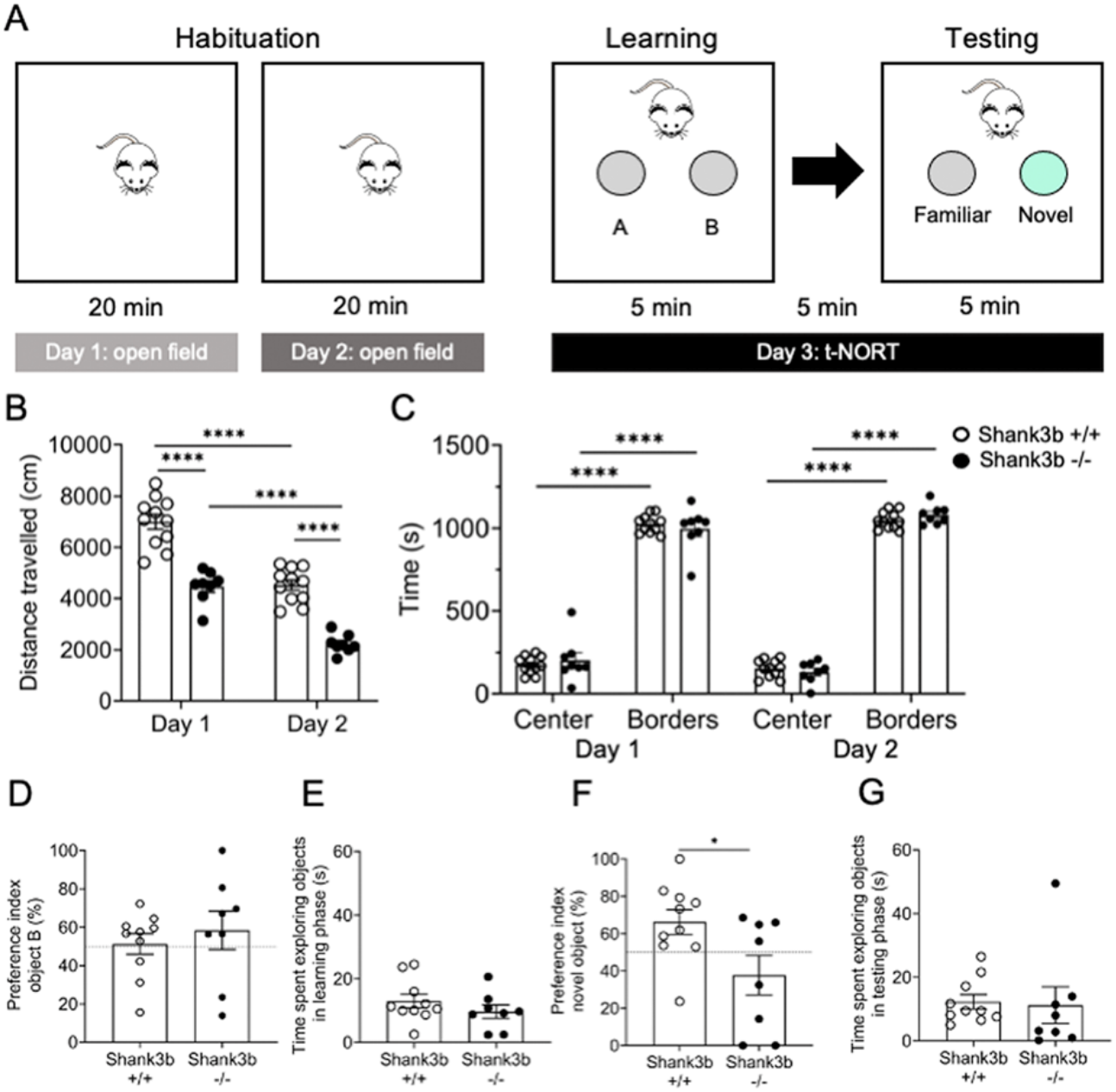
*Shank3b^−/−^* mice exhibit hypo-locomotion and deficits in texture discrimination. (*A*) Open field and texture novel object recognition test (tNORT) experimental design. (*B* and *C*) Quantification of open field performance by *Shank3b^+/+^* and *Shank3b^−/−^* mice. *Shank3b^−/−^* mice travelled a significantly shorter distance over the two days of test, as compared to controls (*B,* ***p < 0.001, ****p < 0.0001, Tukey’s test following two-way ANOVA). Both *Shank3b^+/+^* and *Shank3b^−/−^* mice spent significantly more time in borders as compared to the center of the arena (*C*, ***p < 0.001, ****p < 0.0001, Tukey’s test following three-way ANOVA), but the time spent in center and borders did not significantly differ between genotypes (*C*, p > 0.05, Tukey’s test following three-way ANOVA). (*D*-*G*) Quantification of tNORT performance by *Shank3b^+/+^* and *Shank3b^−/−^* mice. Preference index (%) for object B (*D*) and time spent exploring objects in learning phase (*E*) did not differ between *Shank3b^+/+^* and *Shank3b^−/−^* mice (p > 0.05, Tukey’s test following two-way ANOVA). In the testing phase, *Shank3b^−/−^* mice showed a significantly lower preference index (%) for the novel object, as compared to *Shank3b^+/+^* mice (*F*, *p < 0.05, Tukey’s test following two-way ANOVA), while spending the same time as controls in exploring objects (*G*, p > 0.05, Tukey’s test following two-way ANOVA). In *D* and *F*, the dashed line represents 50% in the preference index, meaning equal preference for both objects. See Materials and Methods for details about preference index calculation. All plots report the mean values ± SEM; each dot represents one animal. Genotypes are as indicated (n=11 *Shank3b^+/+^* and 8 *Shank3b^−/−^* for open field; n=10 *Shank3b^+/+^* and 8 *Shank3b^−/−^* for tNORT; one *Shank3b^+/+^* mouse was excluded from tNORT analysis as it did not interact with objects during the learning phase).

During the tNORT learning phase (Fig. 1*D*) both genotypes did not show any preference for one of the identical textured objects (unpaired t-test, *Shank3b^+/+^* vs. *Shank3b^−/−^*, p=0.5137). During the test phase, control mice spent a significantly larger amount of time exploring the novel object, testifying a preference in the novelty exploration through whiskers. Conversely, *Shank3b^−/−^* mice spent comparable time exploring the novel and the old object, (Fig. 1*F*; unpaired t-test, *Shank3b^+/+^* vs. *Shank3b^−/−^*, p=0.0301; see also *SI Appendix*, Fig. S2). Moreover, the amount of time spent investigating objects during both learning and testing phase did not differ between genotypes (unpaired t-test, *Shank3b^+/+^* vs. *Shank3b^−/−^*; p>0.05; Fig. 1*E* and *G*), indicating that mutant mice did not exhibit an aversion to the objects, and did not avoid object exploration through whiskers. These results suggest that *Shank3b^−/−^* mice display a lack of interest in whisker-mediated exploration, potentially due to reduced whisker-dependent texture discrimination.

### *Shank3b^−/−^* mice display hypo-reactivity to repetitive whisker stimulation

Given their lack of novelty exploration in the tNORT, we reasoned that *Shank3b^−/−^* mice might display impaired ability to interact with their surroundings in a whisker-sensation-dependent context. Thus, we assessed their behavioral responses to whisker stimulation using the whisker nuisance (WN) task (23). *Shank3b^−/−^* and control mice were repetitively stimulated with a wooden stick over 3 sequential trial sessions of 5 min each (Fig. 2*A*). Behaviors were scored according to a 0–10 points scale (*SI Appendix*, Table S1). Both genotypes displayed a comparable response during the pre-stimulation session (sham), showing no anxious behavior when the stick was presented in proximity of the animal’s head, avoiding any contact (Tukey’s post hoc following two-way ANOVA; *Shank3b^+/+^* vs. *Shank3b^−/−^*; p>0.05; Fig. 2*C*). Conversely, *Shank3b^−/−^* mice displayed a significantly lower score across the 3 trials compared with controls (Mann-Whitney test, *Shank3b^+/+^* vs. *Shank3b^−/−^*; p=0.0001; Fig. 2*B*). No differences between sexes were observed within the two genotypes (*SI Appendix*, Fig. S3). Two-way ANOVA revealed a significant effect of genotype and trial (main effect of genotype F_(1, 116)_ = 33.02; p<0.0001; main effect of trial F_(3, 116)_ = 91.00; p<0.0001; Fig. 2*C*). Both genotypes exhibited a significant reduction in the scores assigned to behavioral responses from the first to the third trial, indicating habituation to repetitive whisker stimulation (*SI Appendix*, Fig. S4). However, during the three trials of whisker stimulation, the overall behavioral response of *Shank3b^−/−^* mice was always significantly lower than that of control mice (Tukey’s post hoc following two-way ANOVA; *Shank3b^+/+^* vs. *Shank3b^−/−^*; p<0.01 for trials 1 and 3; p<0,001 for trial 2; Fig. 2*B*). We then analysed the behavioral response of *Shank3b^−/−^* and control mice for each of the five categories used to calculate the WN score. When compared with their controls, *Shank3b^−/−^* mice displayed a significantly lower score for stance, evasion and breathing (Bonferroni’s test following two-way ANOVA, *Shank3b^+/+^* vs. *Shank3b^−/−^*; p=0.0053 for stance, p=0.0030 for evasion, p=0.0001 for breathing; Fig. 2*D*). These findings indicate that *Shank3b^−/−^* mice are less prone to engage in proactive behaviors in response to repetitive whisker stimulation compared to control animals.

**Fig. 2.**
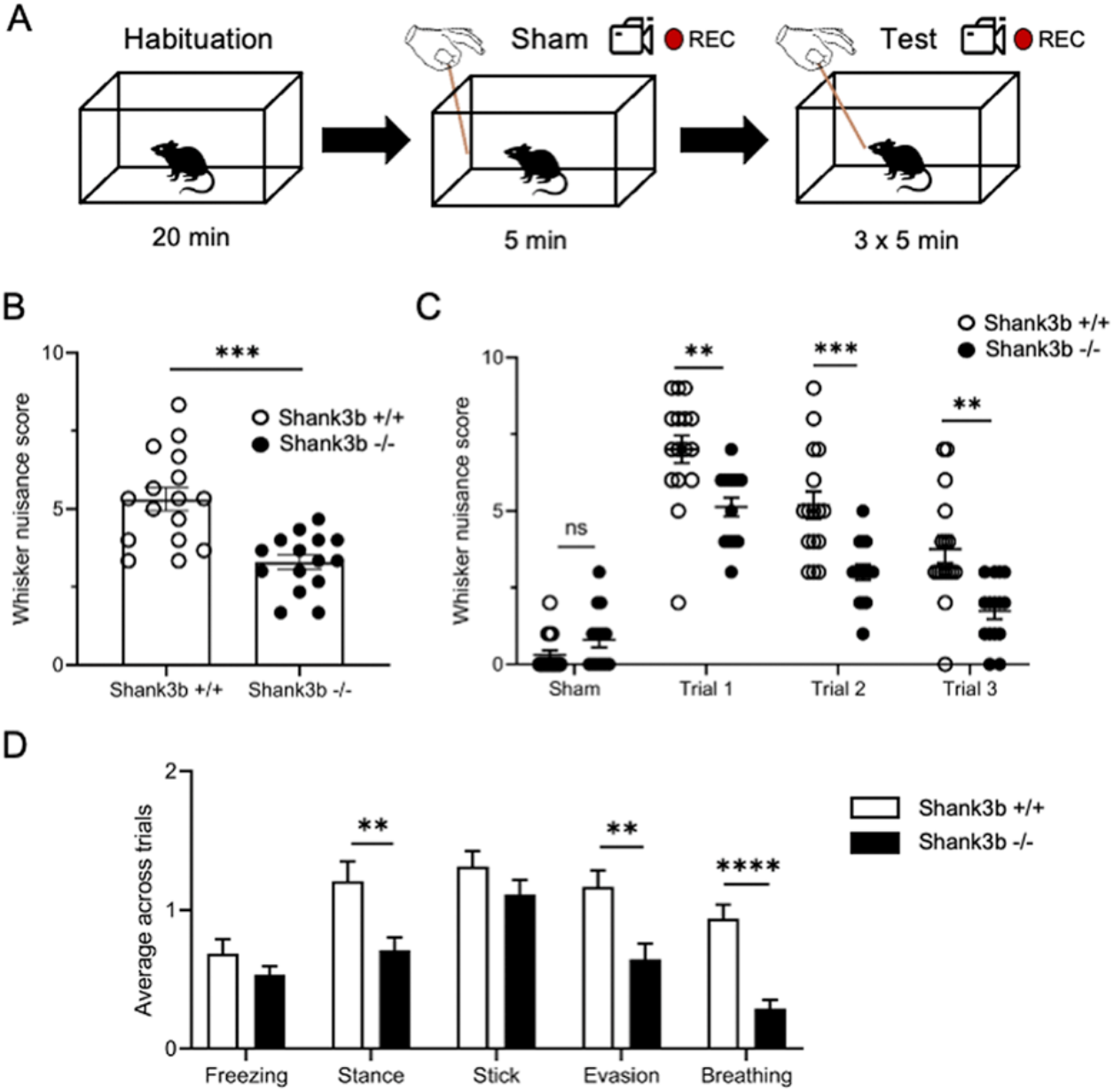
*Shank3b^−/−^* mice are hypo-reactive to repetitive whisker stimulation. (*A*) Whisker nuisance (WN) test experimental design; five different parameters (freezing, stance, response to stick, evasion, and breathing) were monitored across the sham session (no tactile stimulation) and three different experimental trials of repetitive whisker stimulation (see Materials and Methods for WN score calculation). (B-C) Quantification of WN scores of *Shank3b^+/+^* and *Shank3b^−/−^* mice. *Shank3b^−/−^* mice showed a significantly lower total score, as compared to controls (*B*, ***p < 0.001, unpaired t-test; single dots indicate the mean value of WN score for each animal across the three trials). *Shank3b^−/−^* mice scored significantly lower than *Shank3b^+/+−^* mice in all trials, but not during the sham session (*C*, **p < 0.001, ***p < 0.001, Bonferroni’s test following two-way ANOVA; each dot represents one animal). (*D*) Mean scores of *Shank3b^+/+^* and *Shank3b^−/−^* mice for each of the five behavioral parameters monitored across trials (**p < 0.01, ***p < 0.001, Tukey’s test following two-way ANOVA). All plots report the mean values ± SEM. Genotypes are as indicated (n=16 *Shank3b^+/+^* and 15 *Shank3b^−/−^*).

### *Shank3b^−/−^* mice show reduced c-*fos* mRNA expression in S1 and hippocampus following repetitive whisker stimulation

*Shank3b^−/−^* mice sensory hypo-responsiveness in the WN task led us to investigate the neural substrates of aberrant whisker-dependent behaviors in these mutants. To this aim, we used c-*fos* mRNA expression as a molecular proxy for neural activity (23, 24) and quantified *in situ* hybridization signal intensity in multiple regions of *Shank3b^−/−^* and control brains 20 min after either a sham or WN session. Comparable expression of c-*fos* mRNA was observed in both genotypes following the sham session (Fig. 3*A* and *B*). Conversely, significant downregulation of c-*fos* expression was detected in *Shank3b^−/−^* S1, predominantly in layer 4 (Fig. 3*C*; Tukey’s post hoc following two-way ANOVA, *Shank3b^+/+^* vs. *Shank3b^−/−^*; L2/3, p=0.0024; L4, p<0.0001; L5/6, p=0.0065) and hippocampus, predominantly in dentate gyrus (DG) (Fig. 3*D*; Tukey’s post hoc following two-way ANOVA, *Shank3b^+/+^* vs. *Shank3b^−/−^*; CA1, p= 0.0073; CA2, p=0.0495; DG, p=0.0009). No difference was observed in other brain regions (*SI Appendix*, Fig. S5).

**Fig. 3.**
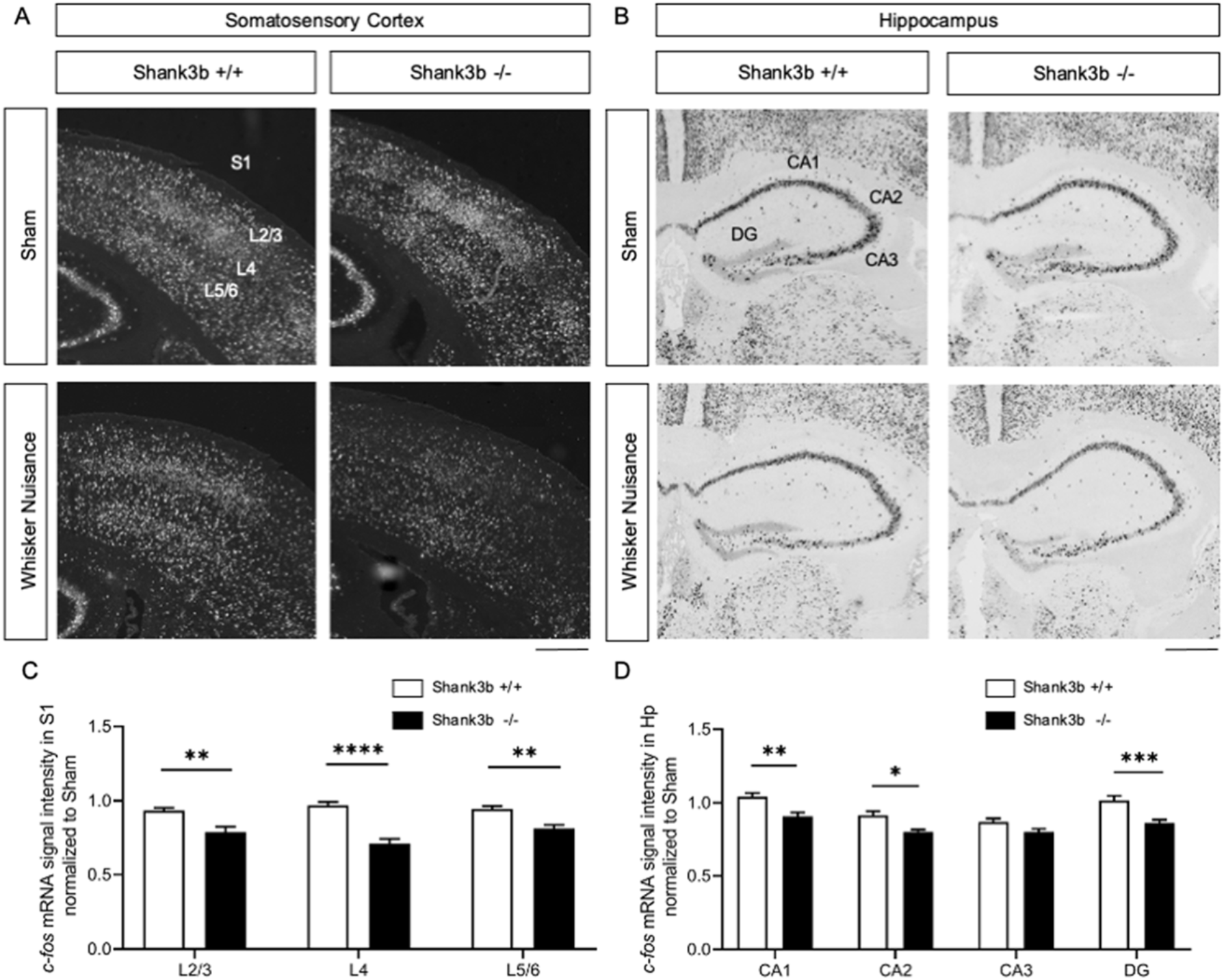
*Shank3b^−/−^* mice show reduced neuronal activation in the primary somatosensory cortex and hippocampus following whisker nuisance test. (*A* and *B*). Representative images of c-*fos* mRNA *in situ* hybridization in the primary somatosensory cortex (S1; white staining) and hippocampus (black staining) of *Shank3b^+/+^* and *Shank3b^−/−^* mice, 20 min following sham or WN test. Scale bars, 500 μM. (*C*) Quantification of c-*fos* mRNA signal intensity in S1, normalized to sham. c-*fos* mRNA downregulation was predominantly detected in layer 4 (Cohen’s d effect-size, Shank3b+/+ vs. Shank3b−/−; L2/3, d=1.21; L4, d=2.15; L5/6, d=1.36). (*D*) Quantification of c-*fos* mRNA signal intensity in the hippocampus, normalized to Sham. c-*fos* mRNA downregulation was predominantly detected in DG (Cohen’s d effect-size, Shank3b+/+ vs. Shank3b−/−; CA1, d=1.20; CA2, d=1.08; CA3, d=0.66; DG, d=1.23). Values are expressed as mean normalized signal intensities ±SEM (see Materials and Methods). *p<0.05, **p<0.01, ***p<0.001, ****p<0.0001, Tukey post-hoc test following two-way-ANOVA (n=22 sections from 6 *Shank3b^+/+^*mice and 16 sections from 4 *Shank3b^−/−^* mice). Genotypes and treatments are as indicated. Abbreviations: CA1/2/3, hippocampal pyramidal cell layers; DG, dentate gyrus; Hp, hippocampus; L2-6, S1 cortical layers; S1, primary somatosensory cortex.

To confirm that the reduced c-*fos* expression observed in the *Shank3b^−/−^* S1 and hippocampus following WN was explicitly due to whisker stimulation, we next investigated c-*fos* mRNA induction in these areas following whisker stimulation in lightly anesthetized *Shank3b^+/+^*and *Shank3b^−/−^* mice. *In situ* hybridization experiments showed a reduction of c-*fos* mRNA mean signal intensity in *Shank3b^−/−^* S1 (Fig. 4*A*) and hippocampus (Fig. 4*C*), compared to controls. Quantification of mean signal intensity confirmed a significantly lower c-*fos* mRNA expression in *Shank3b^−/−^* S1 (unpaired t-test; *Shank3b^+/+^* vs. *Shank3b^−/−^*; p<0.0001; Fig. 4*B*) and hippocampus (unpaired t-test; *Shank3b^+/+^* vs. *Shank3b^−/−^*; p=0.0009; Fig. 4*D*). A significantly lower c-*fos* mRNA expression was also detected in *Shank3b^−/−^* somatosensory thalamus (Vpm, ventral posterior medial nucleus), while no difference was detected in other brain regions analyzed (*SI Appendix*, Fig. S6). These findings indicate that in *Shank3b^−/−^* mice, whisker stimulation results in a marked reduction of c-*fos* mRNA in both S1 and hippocampus. These results point at impaired hippocampal processing of somatosensory signals as a putative large-scale correlate of the observed behavioral phenotype.

**Fig. 4.**
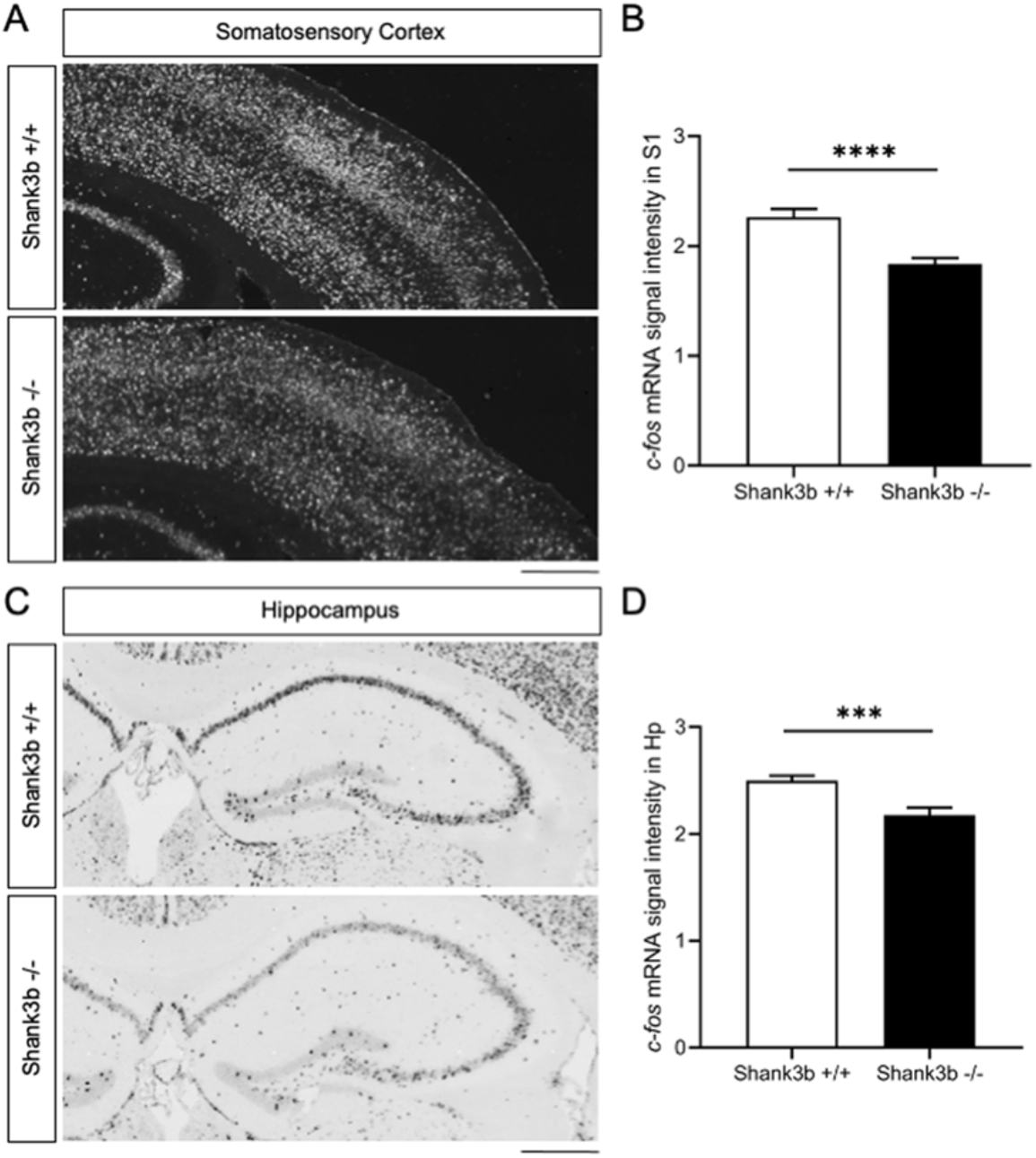
*Shank3b^−/−^* mice show reduced neuronal activation in S1 and hippocampus following whisker stimulation under anesthesia. (*A* and *C*) Representative images of c-*fos* mRNA *in situ* hybridization in the primary somatosensory cortex (A, white staining) and hippocampus (C, dark staining) of *Shank3b^+/+^* and *Shank3b^−/−^* mice, 20 min following whisker stimulation under anesthesia. Scale bars, 500μm. (*B* and *D*) Quantification of c-*fos* mRNA signal intensity in S1 (*B*) and hippocampus (*D*) of *Shank3b^+/+^* and *Shank3b^−/−^* mice. Values are expressed as mean normalized signal intensities ±SEM. ***p<0.001, ****p<0,0001 (unpaired t-test, n=8 sections from 4 animals per genotype).

### Reduced long-range functional connectivity between hippocampus and S1 in *Shank3b^−/−^* mice

We previously showed that loss of *Shank3b* leads to profoundly altered cortico-cortical functional coupling (25). To investigate whether the regional deficits observed upon repetitive whisker stimulation could be linked to similarly impaired hippocampus-S1 functional synchronization, we probed rsfMRI connectivity of the dorsal hippocampus, a brain region characterized by a decreased c-*fos* signal in *Shank3b* mutants (Fig. 3 and 4). Interestingly, rsfMRI mapping revealed markedly reduced functional connectivity between the dorsal hippocampus and S1 (|t| > 2, p<0.05 and FWER cluster-corrected using a cluster threshold of p=0.01, Fig. 5*A*). Quantification of rsfMRI signal in regions of interest confirmed impaired functional connectivity between hippocampus and S1 (t=3.57, p=0.002, Fig. 5*B*), which, like the dorsal hippocampus, we found to be hypo-activated by sensory stimuli in *Shank3b^−/−^* mice (Fig. 3 and 4). These findings suggest that the reduced cortical-hippocampal response observed in *Shank3* mutants might result from a weakened functional coupling between hippocampal and somatosensory areas.

**Fig. 5.**
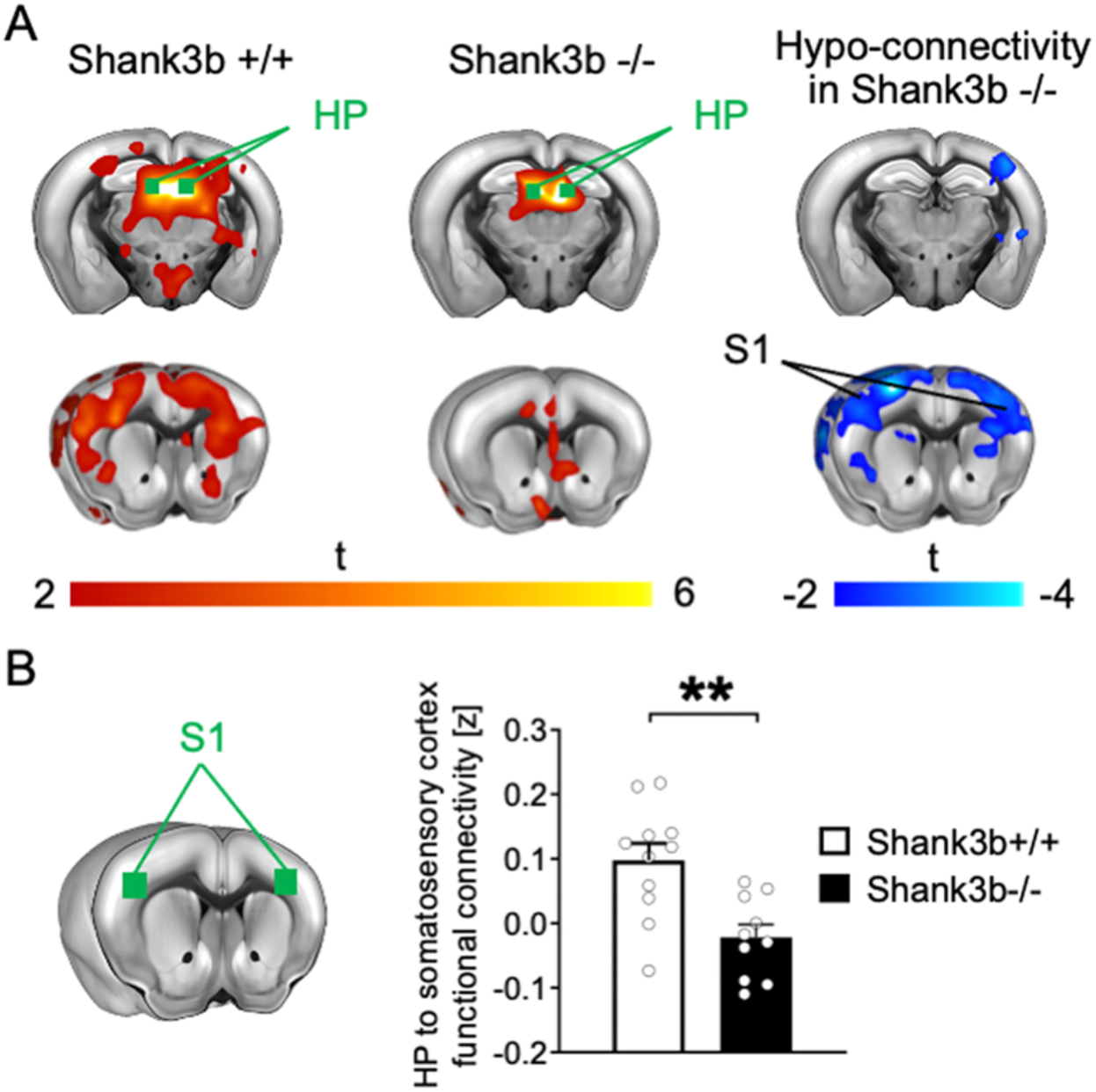
Impaired functional connectivity between hippocampus and primary somatosensory cortex (S1) in *Shank3b^−/−^* mice. (*A*) Seed-based connectivity maps of the dorsal hippocampus in *Shank3b^+/+^* and *Shank3b^−/−^* mice. Red-yellow represents brain regions showing significant rsfMRI functional connectivity with the dorsal hippocampus (HP) in *Shank3b^+/+^* (left) and *Shank3b^−/−^* mice (middle). Seed region is depicted in green. Brain regions showing significantly reduced rsfMRI connectivity in *Shank3b^−/−^* mutants with respect to *Shank3b^+/+^* control littermates are depicted in blue / light blue (right). (*B*) Functional connectivity was also quantified in reference volumes of interest (green) placed in S1. Error bars represent SEM. **p<0.01(unpaired t-test, n= 11 *Shank3b^+/+^* and 10 *Shank3b^−/−^*^;^ each dot represents one animal). Genotypes are as indicated.

## Discussion

In this study, we report that *Shank3b* mutant mice display aberrant whisker-dependent behaviors. This trait is associated with altered response and coupling of circuits involved in sensory processing. Specifically, we found that *Shank3b^−/−^* adult mice showed normal exploration through whiskers but reduced texture discrimination in a whisker-dependent task. *Shank3b^−/−^* mice showed hypo-responsiveness to repetitive whisker stimulation, accompanied by a significantly reduced c-*fos* mRNA expression in S1 and hippocampus. Finally, using rsfMRI, we detected reduced long-range functional connectivity between hippocampus and S1 in mutant mice. We propose that hypo-connectivity and reduced activation of cortico-hippocampal circuits might represent a functional substrate for hypo-reactivity to whisker-dependent cues in *Shank3b^−/−^* mice.

Previous studies investigated tactile discrimination in mouse strains harboring ASD-related mutations, including *Shank3b* mutants (20, 21). In these investigations, tNORT was performed after whisker removal to avoid whisking investigations and promote object exploration via paws’ glabrous skin. The reported results suggest that whisker-independent aberrant texture discrimination is a common trait in ASD mouse models (13, 20, 21). Mice use their whiskers to explore and navigate the surrounding environment (14, 15, 17, 26). Interestingly, whisker-dependent responses are affected in mouse strains bearing mutations in ASD-relevant genes (4, 23, 27–29). Here we sought to investigate the neural substrates of whisker-dependent behaviors in *Shank3b^−/−^* adult mice. We first used a version of tNORT specifically designed to favor whisker-mediated object exploration (see and Materials and Methods) (22). In this task, *Shank3b^−/−^* mice spent significantly less time exploring the novel (differently textured) object than control mice (Fig. 1*F*). Mutant mice spent the same amount of time investigating objects as controls (Fig. 1*E* and *G*). Together, these results indicate that *Shank3b^−/−^* mice do not avoid object interaction through whiskers and that their hypo-locomotor phenotype (Fig. 1*B*) does not prevent whisker-guided exploration. Our findings suggest that *Shank3b^−/−^* mice display a reduced interest in contextual changes in a whisker-dependent task.

We next asked whether *Shank3b^−/−^* mice are less prone to engage in proactive behaviors in response to novelty or threats. We thus used the WN test to study *Shank3b^−/−^* mice behavior in response to active whisker stimulation. This test has been recently used to characterize whisker-dependent behaviors in mice harboring ASD-relevant mutations (23, 28). *Shank3b^−/−^* mice displayed hypo-reactivity in the WN test (Fig. 2*B*), *Shank3b^−/−^* and control mice showed a similar fear response to repetitive whisker stimulation, as testified by comparable freezing behavior scores (Fig. 2*D*). Conversely, mutant mice displayed a significantly reduced avoidance behavior as indicated by their lower score in the evasion and stance categories, accompanied by a diminished breathing score (Fig. 2*D*). We rule out that behavioral hypo-reactivity of *Shank3b^−/−^* mice in the WN test is due to hypo-locomotion. Both genotypes showed a comparable behavioral response during the sham session (Fig. 2*C*), suggesting that anxiety does not affect mice’s performance in the WN test. Finally, we detected no difference in WN score between male and female mice of both genotypes (*SI Appendix*, Fig. S3), suggesting that male and female *Shank3b^−/−^* mice exhibit comparable behavioral responses (21, 30). We conclude that *Shank3b^−/−^* mice display reduced evasive responses to active whisker stimulation. This observation suggests that *Shank3b^−/−^* mice are less prone to engage in proactive behaviors in response to novelty or threats, consistently with their restricted behavioral interactions with a dynamic environment in the tNORT and WN test.

Recent studies showed that repetitive whisker stimulation during the WN test results in the altered expression of the activity marker c-Fos in several brain areas of ASD mouse models (23, 28) mutant mice. Whisker stimulation during the WN test resulted in c-*fos* mRNA downregulation in *Shank3b^−/−^* S1 and hippocampus (Fig. 3), but not other brain areas (*SI Appendix*, Fig. S5). Accordingly, whisker stimulation under anesthesia selectively downregulated c-*fos* mRNA in the same regions of *Shank3b^−/−^* mice (Fig. 4 and *SI Appendix*, Fig. S6). Recent findings showed that reduced firing of cortical GABAergic neurons leads to hyper-reactivity to weak tactile stimulation in a vibrissa motion detection task in *Shank3b^−/−^* mice (29). Accordingly, a reduced expression of the GABAergic marker parvalbumin was detected in *Shank3b^−/−^* brains (21, 31). Our results show that *Shank3b^−/−^* mice are hypo-reactive to repetitive whisker stimulation (Fig. 2) and display reduced c-*fos* mRNA expression in S1 and hippocampus (Fig. 3 and 4). Further studies are needed to clarify whether whisker stimulation results in decreased c-*fos* expression in cortical and hippocampal interneurons in *Shank3b^−/−^* mice. c-*fos* mRNA downregulation observed in *Shank3b^−/−^* hippocampus following repetitive whisker stimulation is consistent with hippocampal synaptic impairment previously shown in *Shank3* mutants (32, 33). Whisker-dependent c-*fos* mRNA downregulation might reflect a diminished hippocampal cell response to repetitive stimulation, suggesting that impaired integration of somatosensory stimuli occurs in the *Shank3b^−/−^* hippocampus

The hippocampus receives inputs from the somatosensory cortex (34), and several studies indicate that many inputs control hippocampal neurons’ responses. CA1 pyramidal cells receive information from the whiskers via the somatosensory thalamus (Vpm) and entorhinal cortex (35). Somatosensory stimulation increases DG granule cell firing, while predominantly inhibitory responses occur in CA1 (36). Hippocampal neurons use afferent somatosensory information to dynamically update spatial maps (35). Accordingly, blockade of tactile transmission by applying lidocaine on the whisker pad decreased the firing rate of hippocampal place cells, resulting in expanding their place fields in the rat (37). These findings indicate that the hippocampus is a key region for the contextual encoding of somatosensory stimuli. Thus, reduced activation of the cortico-hippocampal circuit might affect whisker-guided contextualization and reduce avoidance behavior in *Shank3b^−/−^* mice.

Synchronized activity between S1 and hippocampus is crucial for the formation of cognitive maps. In the rat, coherence between S1 firing and hippocampal activity increases when the animal collects sensory information through whiskers, and such coherence enhances the integration of somatosensory information in the hippocampus (38). In mice, tactile experience enrichment induces c-Fos expression in the hippocampus and improves memory by modulating the activity of DG granule cells that receive sensory information from S1 via the entorhinal cortex (39). Chemogenetic activation of DG neurons receiving tactile stimuli results in memory enhancement, while inactivation of DG or S1-innervated entorhinal neurons has opposite effects (39). Thus, tactile experience modifies cognitive maps by modulating the activity of the S1 to hippocampus pathway, confirming the importance of this circuit in somatosensory information processing. Within this framework, the observed rsfMRI hypo-connectivity between the dorsal hippocampus and S1 might represent a network substrate for the reduced activation of these areas (Fig. 3 and 4) following repetitive whisker stimulation in *Shank3b^−/−^* mice. This observation is of interest also in the light of recent reports of impaired large-scale cortico-cortical connectivity in *Shank*3-deficient mutants (25). This dysconnectivity might also explain why *Shank3b^−/−^* mice exhibit restricted behavioral responses to whisker-dependent novel stimuli or threats. Together, these results point toward S1-hippocampus circuit dysfunction as a basis for somatosensory processing deficits in mice harboring ASD-relevant mutations. Further studies are needed to extend these findings to ASD.

## Materials and Methods

### Animals

Animal research protocols were reviewed and approved by the University of Trento animal care committee and Italian Ministry of Health, in accordance with the Italian law (DL 26/2014, EU 63/2010). Animals were housed in a 12 h light/dark cycle with food and water available *ad libitum*. All surgical procedures were performed under anesthesia and all efforts were made to minimize suffering. *Shank3b* mutants were crossed at least five times into a C57BL/6 background. Heterozygous mating (*Shank3b^+/−^* x *Shank3b^+/−^*) was used to generate the *Shank3b^+/+^* and *Shank3b^−/−^* littermates used in this study. PCR genotyping was performed according to the protocol available on The Jackson Laboratory website (www.jax.org). A total of 85 age-matched adult littermates (42 *Shank3b^+/+^* and 37 *Shank3b^−/−^*; 3–6 months old; weight = 25–35 g) of both sexes were used. Twenty-one mice (11 *Shank3b^+/+^* and 10 *Shank3b^−/−^*) were used for fMRI experiments and 50 mice (27 *Shank3b^+/+^* and 23 *Shank3b^−/−^*) were used for behavioral testing. A subset of animals subjected to the WN test (6 *Shank3b^+/+^* and 4 *Shank3b^−/−^*) was used for c-*fos* mRNA *in situ* hybridization. An additional group of 3 *Shank3b^+/+^* and 3 *Shank3b^−/−^* mice received only sham stimulation and were used as controls for *in situ* hybridization experiments (no difference in WN score was detected between genotypes in this additional group of animals). Finally, 4 mice per genotype were used for the c-*fos* mRNA study following whisker stimulation under anesthesia. Previous studies showed that similar group sizes are sufficient to obtain statistically significant results in fMRI (25, 40, 41), behavioral, and *in situ* hybridization studies (23, 42, 43). All experiments were performed blind to genotype. Animals were assigned a numerical code by an operator who did not take part in the experiments and codes were associated to genotypes only at the moment of data analysis.

### Open field test

Mice were habituated to the testing arena prior to the texture discrimination assessment, during two consecutive days (Fig. 1*A*). During these two sessions, each animal was placed in empty arena (40 cm × 40 cm × 40 cm) and allowed to freely explore for 20 min. The walls of the arena were smooth and grey colored. Sessions were recorded and mice were automatically tracked using EthoVisionXT (Noldus). Distance travelled and time spent in center/borders were analyzed for both days of test.

### Textured novel object recognition test (tNORT)

Whisker-mediated texture discrimination was assed as described (22), with minor modifications. tNORT was performed in the same arena used for open field testing (Fig. 1*A*). On the third day, cylinder-shaped objects (1.5 cm radius base x 12 cm height) covered in garnet sandpaper were placed into the arena test. The grit (G) of the object (i.e., how rough the sandpaper is) was chosen according to Wu et al. 2013 (22), to favor whisker interaction. A 120 G sandpaper (very fine) was used for the familiar object, whereas 40 G sandpaper (very coarse) was used for the novel object. Many identical objects were created for each grit of sandpaper used in this study to avoid repetitive use of the same object across the testing period. This minimized the possibility that mice recognized one particular object using olfactory cues. In the first session (learning phase), mice were placed in the testing arena facing away from two identically textured objects (object A and object B; 120 G) and allowed to freely explore the objects for 5 min. This short time was selected to favor the investigation through whiskers. The textured objects were placed in the center of the arena, equidistant to each other and the walls. Mice were then removed and held in a separate transport cage for 5 minutes. This short time was selected to minimize hippocampal mediated learning (22). Prior to the start of the second session, the two objects in the arena were replaced with a third, identically textured object (familiar, 120 G) and a new object with different texture (novel, 40 G). The position of the novel versus the familiar object was counterbalanced and pseudorandomized. Mice were then placed back into the arena for the second session (testing phase) and allowed to explore for 5 min. Since mice have an innate preference for novel stimuli, an animal that can discriminate between the textures of the objects spends more time investigating the novel textured object, whereas an animal that cannot discriminate between the textures is expected to investigate the objects equally. The testing arena was cleaned with 70% ethanol between sessions and between animals to mask olfactory cues. The amount of time mice spent actively investigating each of the objects was assessed during both learning and testing phases. Investigation was defined as directing the nose towards the object with a distance of less than 2 cm from the nose to the object or touching the nose to the object. Resting, grooming, and digging next to, or sitting on, the object was not considered as investigation. If an animal did not interact with both objects during the learning phase, it was excluded from tNORT analysis. The activity of the mice during the learning and testing phase was recorded with a video camera centered above the arena and automatically tracked using EthoVisionXT (Noldus). The performance of the mice in the tNORT was expressed by the preference index. The preference index is the ratio of the amount of time spent exploring any one of the two objects in the learning phase or the novel one in the testing phase over the total time spent exploring both objects, expressed as percentage [i.e., A/(B + A) × 100 in the learning session and novel/(familiar + novel) × 100 in the testing session].

### Whisker nuisance (WN) test

WN test was performed as previously described (23, 44). Briefly, animals were allowed to habituate for 30 minutes to a novel empty cage (experimental cage) for 2 days before the test. To facilitate the habituation to the novel environment, home-cage bedding was placed overnight in the experimental cage and removed right before the introduction of the mouse. The proper testing session lasted approximately 1 hour per mouse, divided into a 35 min pre-test (30 min of environment habituation and 5 minutes of experimenter habituation) followed by a 20 min test. The 20 min test was split into 4 sessions of 5 min each. In the first session (“sham”), the wooden stick was introduced in the cage avoiding any tactile contact with the mouse whiskers or body. Following sham stimulation, mice underwent three consecutive stimulation sessions (trials 1–3, with 1 min of inter-trial interval), each consisting of continuous bilateral stimulation of the whiskers with a wooden stick. The predominant behavioral response during the four test sessions was scored over a 10-point scale (see also *SI Appendix*, Table S1) (23, 44), which represents a modified version of the WN test originally developed to assess sensory behaviors in rats (45). Scoring criteria were divided into five categories (freezing, stance, response to stick presentation, evasion, breathing). Normal behavioral responses to stimulation were assigned a 0 value, whereas meaningful abnormal behavioral responses (i.e., for the vast majority of the observation period) were assigned a value of 2. The maximum WN score is 10. High scores (8–10) indicate over-responsiveness to the stimulation, while low scores (0–3) indicate hypo-responsiveness (*SI Appendix*, Table S1). Scoring was performed by two independent experimenters blind of genotype, and no variability in scoring was observed.

### Whisker stimulation under anesthesia

Mice were anesthetized with an intraperitoneal injection of urethane (20% solution in sterile double-distilled water, 1.6 g/kg body weight) and head-fixed on a stereotaxic apparatus. Urethane anesthesia was chosen as it preserves whisker-dependent activity in the somatosensory cortex (46). WS protocol consisted in 3 consecutive sessions (5 minutes each, with 1 min intervals) of continuous touch of the whiskers with a wooden stick (bilateral stimulation), thus reproducing the stimulation protocol used in the WN test (23).

### c-*fos* mRNA *in Situ* Hybridization

Mice were killed 20 min after the end of either sham, WN or WS, and brains were rapidly frozen on dry ice. Coronal cryostat sections (20 μm thick) were fixed in 4% paraformaldehyde and processed for non-radioactive *in situ* hybridization (47) using a digoxigenin-labelled c-*fos* riboprobe. Signal was detected by alkaline phosphatase-conjugated anti-digoxigenin antibody followed by alkaline phosphatase staining. Brain slices from different experimental paradigms (sham, WN, and WS) were processed together for c-*fos* expression in order to exclude possible batch effects. Sense riboprobes, used as negative control, revealed to detectable signal (data not shown). Brain areas were identified according to the Allen Mouse Brain Atlas (https://mouse.brain-map.org). Digital images from 2–4 sections per animal were acquired at the level of the S1/dorsal hippocampus using a Zeiss AxioImager II microscope at 10× primary magnification.

### Behavioral and in situ hybridization data analysis

Statistical analyses of behavioral and *in situ* hybridization data were performed with GraphPad Prism 8.0 software, with the level of significance set at p<0.05. For behavioral experiments, statistical analysis was performed by unpaired t-test or two/three-way ANOVA followed by Tukey’s or Bonferroni’s post-hoc multiple comparisons, as appropriate. To quantify c-*fos* mRNA signal intensity, acquired images were converted to 8-bit (grey-scale), inverted, and analyzed using the ImageJ software (https://imagej.net/Downloads). Mean signal intensity was measured in different counting areas drawn to identify S1 cortical layers, hippocampal subfields and other areas of interest. Mean signal intensity was divided by the background calculated in layer 1. Since behavioral responses did not differ between the two genotypes during the sham session (Fig. 2*C*), c-*fos* mRNA signal intensity in WN animals was normalized to the mean signal intensity value of sham-treated animals of the same genotype. Statistical analysis was performed by unpaired t-test or two-way ANOVA followed by Tukey’s for post-hoc multiple comparisons, as appropriate.

### Resting-State Functional MRI (rsfMRI)

#### Data collection

Resting-state functional MRI (rsfMRI) time series acquisition in adult male *Shank3b^+/+^* (n=11) and *Shank3b^−/−^* (n=10) mice has been previously described (25). All functional connectivity analyses reported here were carried out on the rsfMRI scans acquired for the Pagani et al. 2019 study (25). The protocol for animal preparation employed was detailed in our previous studies (25, 48–50). Briefly, animals were anaesthetized with isoflurane (5% induction), intubated and artificially ventilated (2% maintenance). After surgery, isoflurane was discontinued and replaced with halothane (0.7%). Recordings started 45 min after isoflurane cessation. Functional scans were acquired with a 7T MRI scanner (Bruker Biospin, Milan, Italy) using a 72-mm birdcage transmit coil and a 4-channel solenoid coil for signal reception. For each animal, *in-vivo* anatomical images were acquired with a fast spin echo sequence (repetition time [TR] = 5500 ms, echo time [TE] = 60 ms, matrix 192 × 192, field of view 2 × 2 cm, 24 coronal slices, slice thickness 500 μm). Co-centerd single-shot BOLD rsfMRI time series were acquired using an echo planar imaging (EPI) sequence with the following parameters: TR/TE 1200/15 ms, flip angle 30°, matrix 100 × 100, field of view 2 × 2 cm, 24 coronal slices, slice thickness 500 μm for 500 volumes.

#### Functional connectivity data analysis

Before mapping rsfMRI connectivity, raw data was preprocessed and denoised as previously described (41, 49, 51). First, the initial 50 volumes were removed to allow for T1 equilibration effects. Time series were then despiked, motion corrected and registered to a common reference template. Motion traces of head realignment parameters and mean ventricular signal were then used as nuisance covariates and regressed out from each time course. Before functional connectivity mapping, all time series underwent also band-pass filtering (0.01–0.1 Hz) and spatial smoothing (FWHM = 0.6 mm). Target regions of long-range connectivity alterations in *Shank3b^−/−^* mice were mapped using seed-based analysis. Specifically, bilateral seeds of 3×3×1 voxels were placed in the dorsal hippocampus of *Shank3b^+/+^* and *Shank3b^−/−^* mice to probe impaired functional connectivity between this region and the rest of the brain. The location of the bilateral seeds employed for mapping is indicated in Fig. 5*A*. Functional connectivity was measured with Pearson’s correlation and *r*-scores were transformed to *z*-scores using Fisher’s *r*-to-*z* transform before statistics. Voxel-wise intergroup differences for seed-based mapping were assessed using a 2-tailed Student’s t-test (|*t*| > 2, p<0.05) and family-wise error (FWER) cluster-corrected using a cluster threshold of p=0.01. To quantify rsfMRI alterations we also carried out functional connectivity measures in cubic regions of interest (3×3×1 voxels). The statistical significance of these region-wise intergroup effects was quantified using a 2-tailed Student’s *t*-test (|*t*| > 2, p<0.05).

## Acknowledgments

YB dedicates this study to the memory of Paolo Sassone-Corsi, a great scientist and a dear friend, who unexpectedly left us on July 22, 2020. The c-*fos* cDNA used to prepare riboprobes for *in situ* hybridization experiments was a kind gift of Paolo’s laboratory in 1997. The authors thank Gabriele Chelini (University of Mississippi Medical Center, Jackson, MS) for his valuable comments on the manuscript. The authors thank the technical and administrative staff of CIMeC, CIBIO, and IIT for assistance. This work was supported by the Strategic Project TRAIN - Trentino Autism Initiative (https://projects.unitn.it/train/index.html) from the University of Trento (grant 2018-2023 to YB). LB is a recipient of a PhD fellowship from the University of Trento. LP is supported by a postdoctoral fellowship from the Umberto Veronesi Foundation (Milan, Italy). GP is supported by the Brain and Behavior Research Foundation (NARSAD Young Investigator Grant; ID: 26617) and the University of Trento (Starting Grant for Young Researchers). A.Go. was funded by the Simons Foundation (SFARI 400101), Brain and Behavior Foundation (NARSAD - National Alliance for Research on Schizophrenia and Depression), the European Research Council (ERC-DISCONN, GA802371), the NIH (1R21MH116473-01A1) and the Telethon Foundation (GGP19177).

