## Supplementary material for "Somatosensory processing deficits and altered cortico-hippocampal connectivity in *Shank3b^−/−^* mice": SI Appendix

### Supplementary information

#### Supplementary Figure 1

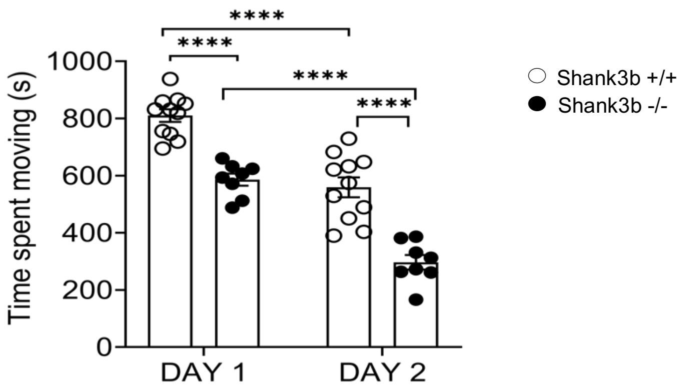


**Fig. S1.** *Shank3b$^{-/-}$* mice spent a significantly shorter time moving in the arena over the two days of test, as compared to controls (*Shank3b$^{-/-}$* vs. *Shank3b$^{+/+}$*, ****p < 0.0001, Tukey's test following two-way ANOVA). Both genotypes showed a significantly reduced time spent moving between day 1 and day 2 (day 1 vs. day 2 within each genotype, ****p < 0.0001, Tukey's test following two-way ANOVA). Bars report mean values ± SEM; each dot represents one animal. Genotypes are as indicated (n=11 Shank3b+/+ and 8 Shank3b-/-)

#### Supplementary Figure 2

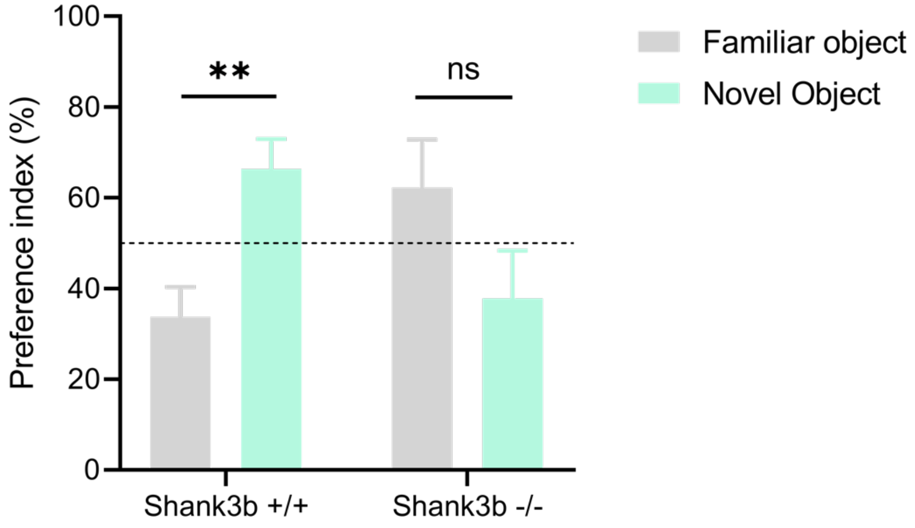


**Fig. S2.** *Shank3b$^{+/+}$* but not *Shank3b$^{-/-}$* mice show a preference for the novel textured object. Values are expressed as mean values (±SEM). Dashed line represents 50% in the preference index (see Fig. 1F), meaning equal preference for both objects. **p<0.01. Multiple t-test, familiar object vs. novel object within *Shank3b$^{+/+}$* and *Shank3b$^{-/-}$* mice (n=10 *Shank3b$^{+/+}$* and 8 *Shank3b$^{-/-}$*).

**Supplementary Figure 3**

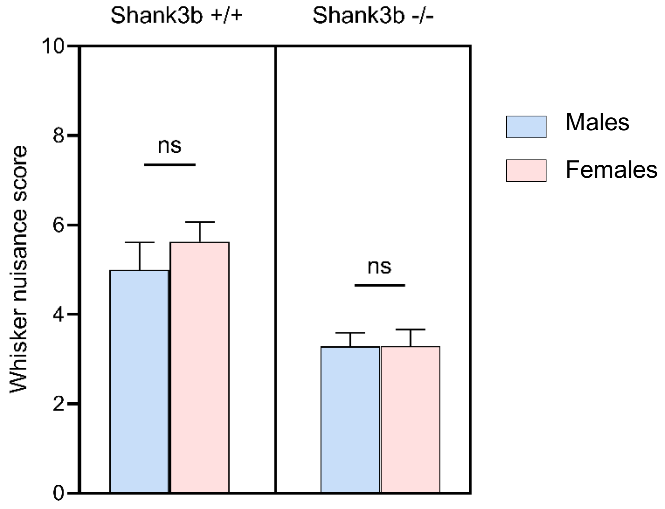


**Fig. S3.** WN total scores did not differ between male and female mice within *Shank3b*$^{+/+}$ and *Shank3b*$^{-/-}$ mice (2-way ANOVA, main effect of sex $F_{(1, 27)}$ = 0,4825, p=0,4932; n = 8 males and 8 females *Shank3b*$^{+/+}$, 7 males and 8 females *Shank3b*$^{-/-}$). Bars report mean values ± SEM. Genotypes and sexes are as indicated.

**Supplementary Figure 4**

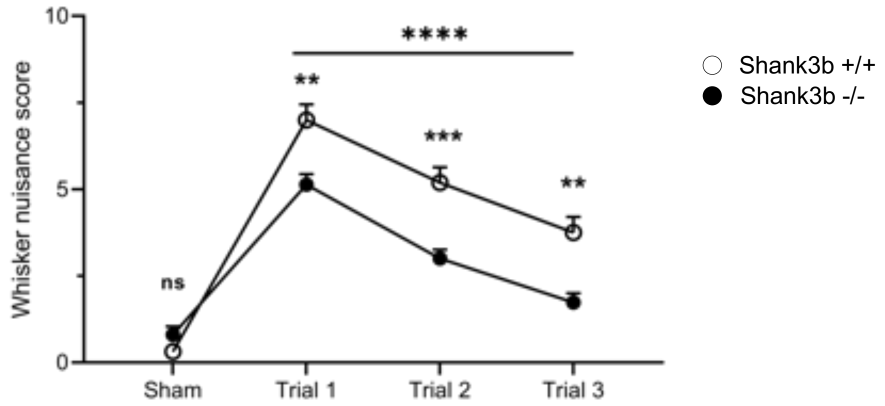


**Fig. S4.** Both *Shank3b*$^{+/+}$ and *Shank3b*$^{-/-}$ mice showed habituation to repetitive whisker stimulation from the first to the third trial of the WN test (Tukey's post hoc following two-way ANOVA; trial 1 vs trial 3 within each genotype ****p< 0.0001). *Shank3b*$^{-/-}$ mice scored significantly lower than *Shank3b*$^{+/+}$ mice in all trials, but not during the sham session (**p < 0.001, ***p < 0.001, Bonferroni's test following two-way ANOVA; see also Fig. 2C). The graph shows WN scores (mean values ± SEM) assigned to *Shank3b*$^{+/+}$ and *Shank3b*$^{-/-}$ mice during sham session and trials. Genotypes are as indicated.

**Supplementary Figure 5**

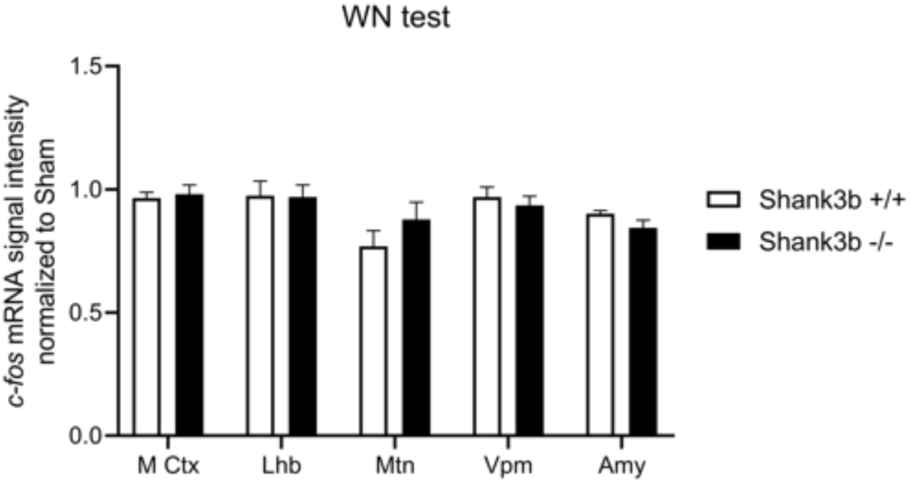


**Fig. S5.** *Shank3b$^{+/+}$* and *Shank3b$^{-/-}$* mice did not show any difference in c-*fos* mRNA signal intensity following WN test in the brain regions analyzed. Values are expressed as mean signal intensities (±SEM), normalized to sham (unpaired t-test, p>0.05 for all comparisons; n=22 sections from 6 *Shank3b$^{+/+}$* mice and 16 sections from 4 *Shank3b$^{-/-}$* mice). Abbreviations: M Ctx, motor cortex; Lhb, lateral habenula; Mtn, medial thalamic nuclei; Vpm, ventral postero-medial nucleus of thalamus; Amy, amygdala.

**Supplementary Figure 6**

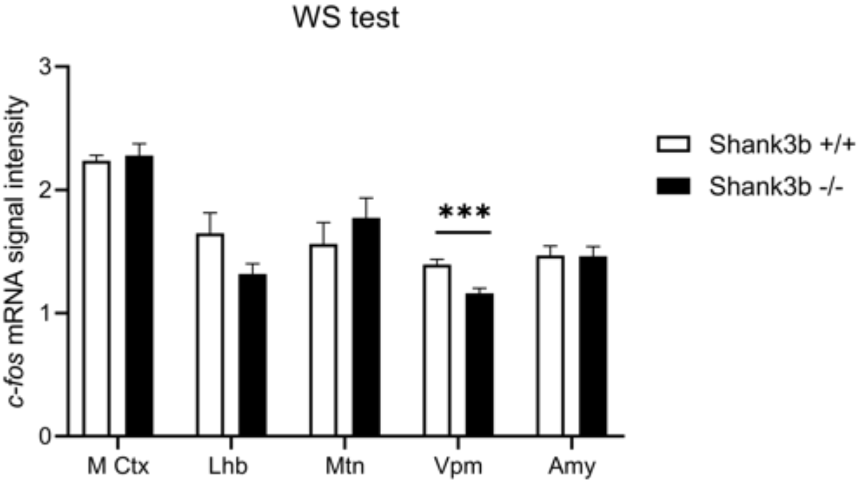


**Fig. S6.** *Shank3b$^{-/-}$* mice did not show reduced c-*fos* mRNA expression in several brain regions following whisker stimulation (WS) under anesthesia. c-*fos* mRNA dowregulation was detected in the ventral postero-medial nucleus of the thalamus (Vpm), which receives afferents from whiskers via brainstem nuclei and projects to S1 layer 4 (Petersen, 2007). Values are expressed as mean normalized signal intensities (±SEM) (unpaired t-test, ***p<0.001 in Vpm, p>0.05 for all other brain areas; n=8 sections from 4 animals per genotype). Abbreviations: M Ctx, motor cortex; Lhb, lateral habenula; Mtn, medial thalamic nuclei; Vpm, ventral postero-medial nucleus of thalamus; Amy, amygdala.

**Supplementary Table 1**

| **Behaviors** | **Whisker Nuisance Score** | | |
|---|---|---|---|
| | 0 | 1 | 2 |
| Fearful behavior | curious, insensible | ambivalent | freezing |
| Stance | skyward head, relaxed | ambivalent | guarded |
| Breathing | normal | ambivalent | hyperventilated |
| Response to stick | interested, ignore | ambivalent | attacking, startle |
| Evasion | explorative bahavior | ambivalent | run away, avoiding |
| **Total score** | **0 to 3: curious, restful** | **4 to 7: annoyed, bothered** | **8 to 10: scared, worried** |

**Table S1. WN scoring.** The predominant behavioral response during each test session (sham and trials 1–3) was scored over a 10-point scale. Normal behavioral responses were assigned a 0 value, while abnormal behavioral responses (i.e., for the vast majority of the observation period) were assigned a value of 2. The maximum WN score is 10. High scores (8–10) indicate over-responsiveness to stimulation, while low scores (0–3) indicate hypo-responsiveness.
